# Col6a1^+^/CD201^+^ mesenchymal cells regulate intestinal morphogenesis and homeostasis

**DOI:** 10.1101/2021.02.16.431453

**Authors:** Maria-Theodora Melissari, Ana Henriques, Christos Tzaferis, Alejandro Prados, Michalis E. Sarris, Panagiotis Chouvardas, Sofia Grammenoudi, George Kollias, Vasiliki Koliaraki

**Affiliations:** Institute for Fundamental Biomedical Research, Biomedical Sciences Research Center (B.S.R.C.) “Alexander Fleming”, Vari, 16672, Greece; Institute for Bioinnovation, Biomedical Sciences Research Center (B.S.R.C.) “Alexander Fleming”, Vari, 16672, Greece; Department of Medical Oncology, Inselspital, Bern University Hospital, University of Bern, Switzerland; Department for BioMedical Research, University of Bern, Bern, Switzerland; Department of Physiology, Medical School, National and Kapodistrian University of Athens, Athens, 11527, Greece

**Author notes:** M.T.M and A.H. contributed equally to this work. Institute for Research in Biomedicine (IRB Barcelona), The Barcelona Institute of Science and Technology, Barcelona, Spain. correspondence: Vasiliki Koliaraki.

## Abstract

Intestinal mesenchymal cells encompass multiple subsets, whose origins, functions, and pathophysiological importance are still not clear. Here, we used the *Col6a1^Cre^* mouse, which targets telocytes and perivascular cells that can be further distinguished by the combination of the CD201, PDGFRα and αSMA markers. Developmental studies revealed that the *Col6a1^Cre^* mouse also targets mesenchymal aggregates that are crucial for intestinal morphogenesis and patterning, suggesting an ontogenic relationship between them and homeostatic telocytes. Cell depletion experiments in adulthood showed that Col6a1^+^/CD201^+^ mesenchymal cells regulate homeostatic enteroendocrine cell differentiation and epithelial proliferation. During acute colitis, they expressed an inflammatory and extracellular matrix remodeling gene signature, but they also retained their properties and topology. Notably, both in homeostasis and tissue regeneration, they were dispensable for normal organ architecture, while CD34^+^ mesenchymal cells expanded, localised at the top of the crypts and showed increased expression of villous-associated morphogenetic factors, providing thus evidence for the plasticity potential of distinct mesenchymal populations in the intestine. Our results provide a comprehensive analysis of the identities, origin, and functional significance of distinct mesenchymal populations in the intestine.

## Introduction

The mammalian intestine is characterized by a unique architecture, which ensures both efficient nutrient and water absorption and rapid self-renewal of the intestinal epithelium. Self-renewal is mediated by Lgr5^+^ multi-potent crypt-base stem cells (CBCs) that progressively give rise to transit amplifying (TA) progenitor cells and differentiated epithelial cell populations with specific absorptive or secretive functions [1, 2]. The tight regulation of this architectural organization is mediated by a gradient of factors produced both by epithelial and stromal cells. Among stromal cells, intestinal mesenchymal cells (IMCs) have emerged as an important cell type for the development and homeostasis of the intestine, by providing both structural support and regulatory elements [3]. Of particular interest is their contribution to the maintenance of the stem cell niche via the production of soluble mediators [4]. Notably, in the absence of epithelial Wnts, production of stromal Wnts is sufficient for the maintenance of epithelial proliferation, while depletion of Foxl1^+^ telocytes or Grem1^+^ trophocytes, which produce niche-supporting factors led to the disruption of the intestinal structure [5–7]. Additionally, BMP production by villous mesenchymal cells inhibits proliferation and favors epithelial cell differentiation, which ensures epithelial homeostasis and supports specialized epithelial functions [8, 9]. The production of such signals is believed to be induced and maintained via the reciprocal communication with epithelial cells. This has been convincingly shown during embryonic development, where PDGF and Hh proteins secreted from the endodermal epithelium act on the underlying mesenchyme to induce the formation of PDGFRα^+^ aggregates, which express BMPs and regulate vilification of the intestine [10].

Beyond their homeostatic functions, intestinal fibroblasts contribute significantly to tissue damage and inflammation. During such conditions, resident fibroblastic cells are activated to produce pro-inflammatory cytokines and chemokines, angiogenic factors, as well as extracellular matrix (ECM) components and remodeling enzymes to facilitate acute inflammatory responses. Deregulation of these processes or chronic injury can lead to chronic inflammatory disorders and fibrosis [11]. Indeed, recent data point to an important role of the microenvironment in shaping cellular programs and driving epithelial regeneration, as fibroblast-specific deletion of the Wnt regulator porcupine (Porcn) or R-spondin 3 led to severely impaired intestinal regeneration [12, 13].

Until recently, mesenchymal cells in the intestine, although known to constitute a group of cell types, were frequently studied as one, mainly due to difficulties in marking, isolating and genetically targeting specific populations. However, recent single-cell transcriptomic analyses of the normal mouse and human intestine revealed the underappreciated extent of mesenchymal heterogeneity and identified several fibroblast subsets with distinct expression profiles and functions [4, 7, 14–16]. However, their origin and spatial organization, as well as the mechanisms through which distinct subsets coordinate signaling gradients along the crypt-villus axis in homeostasis and disease remain elusive. The use of specific markers and Cre-expressing mouse lines have begun to provide such information and are crucial for addressing these issues. Examples include CD34, Foxl1, PDGFRα and Gli1 positive IMCs, which act as critical regulators of the intestinal stem cell niche through their production of Wnts, R-spondins and Gremlin 1, although they are not strictly restricted to single cell types [6, 7, 12, 13, 17, 18]. Lgr5^+^ villous tip telocytes were also recently shown to regulate epithelial villus tip gene expression programs [19]. Single-cell analysis of the inflamed intestine has further highlighted the prominent pro-inflammatory activation of fibroblasts [14, 15, 20]. Notably, activated fibroblasts along with immune cells were associated with resistance to anti-TNF therapy, indicating their potential utility in patient diagnosis, stratification and therapeutic decisions [15, 20].

We have previously shown that the *Col6a1^Cre^* transgenic mouse targets a fraction of mesenchymal cells in the intestine and that NFκΒ signaling in this subset uniquely protects against colitis and colitis-associated cancer [21]. In this study, transcriptomic, imaging and functional analysis of the cells targeted by the *Col6a1^Cre^* mouse revealed preferential targeting of colonic PDGFRα^hi^ telocytes/subepithelial myofibroblasts (SEMFs) and perivascular cells, which can be further described by the use of specific markers. Using reporter mice, confocal imaging and cell depletion approaches we have further identified their relationship with mesenchymal clusters during embryogenesis, as well as their functional significance in development and adulthood. Notably, both in normal conditions and during tissue regeneration, CD34^+^ IMCs proliferated, and adopted different topologies and gene expression profiles, following the depletion of colonic *Col6a1^Cre+^* mesenchymal cells, revealing the plasticity of IMCs towards organ homeostasis.

## Results

### The *Col6α1^Cre^* mouse targets predominantly CD34^−^ mesenchymal cells in the mouse colon

We have shown in the past that the *Col6a1^Cre^* transgenic mouse targets a fraction of mesenchymal cells in the intestine [21]. To define its specificity for distinct mesenchymal subsets, we crossed *Col6a1^Cre^* mice with a TdTomato-to-GFP replacement (mTmG) reporter strain (*Col6a1^mTm^*^G^) and after exclusion of immune, endothelial, erythroid and epithelial cells using the lineage negative (Lin^−^) markers CD45, CD31, Ter119 and CD326 (EpCAM), we isolated *Col6a1*-GFP^+^ and *Col6a1*- GFP^−^ IMCs by FACS sorting (Figure 1A). As previously reported, the *Col6a1^Cre^* mouse does not target Lin^+^ cells [21]. We then performed 3’ mRNA sequencing of the Lin^−^GFP^+^ IMCs (GC), Lin^−^Tomato^+^ IMCs (TC) and the initial unsorted cells (UC). Gene Ontology (GO) analysis of the upregulated genes in the GC versus the TC and/or UC cells revealed enrichment in biological processes related to epithelial cell proliferation and differentiation, as well as regulation of vasoconstriction and blood pressure (Figure 1B). Enriched genes associated with epithelial differentiation included Bmps (*Bmp3*, *Bmp7*, *Bmp2*, *Bmp5*), *Wnt5a,* the Wnt inhibitor *Wif1* and genes related to the differentiation of epithelial cells (e.g. *Fgf9*) [8]. Conversely, the TC population expressed genes associated with the maintenance of the stem cell niche, such as *Grem1*, *Wnt2* and *Nog* [7, 17] (Figure 1B). These results suggest that *Col6a1^Cre+^* cells have distinct homeostatic functions, potentially associated with different topologies along the colonic crypt length.

**Figure 1.**
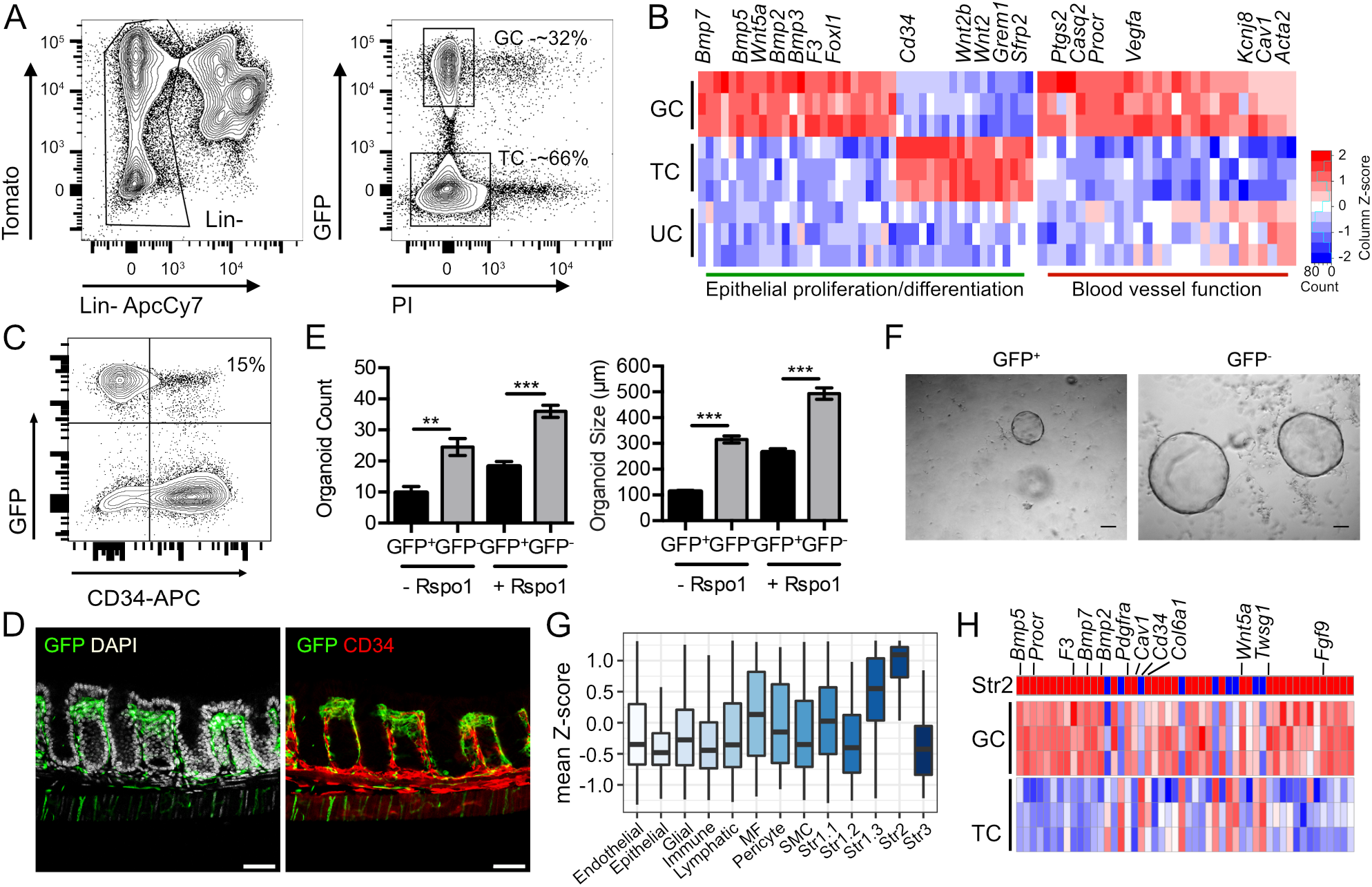
The *Col6a1^Cre^* mouse targets predominantly CD34^−^ IMCs in the mouse colon. A) FACS sorting strategy for the isolation of *Col6a1^Cre^*-GFP^+^ (GC) and *Col6a1^Cre^*-GFP^−^ (TC) mesenchymal cells from the colon. Single cell preparations from the colon (UC) were stained for the Lin^+^ markers CD45, EpCAM, CD31 and Ter119 and Propidium Iodide (PI) for dead cell exclusion. 3 samples from 4-5 mice each were subsequently analyzed. B) Heatmap of differentially expressed genes in GC vs TC and UC samples, corresponding to GO terms related to epithelial proliferation/differentiation and blood vessel regulation. Log2 transformed normalized read counts of genes are shown. Read counts are scaled per column, red denotes high expression and blue low expression values. C) Representative FACS analysis of CD34 expression in Lin-cells in the colon of *Col6a1^mTmG^* mice (n = 4-5 mice). D) Immunohistochemical analysis for CD34 expression in the colon of *Col6a1^mTmG^* mice (n = 9-10 mice, Scale bar: 50 μm). E) Total number and size of intestinal structures after 3 days of co-culture with sorted *Col6a1*-GFP^+^ and GFP^−^ colonic IMCs, with and without R-Spondin 1, respectively. Data represents mean ± SEM from one of four experiments performed in quadruplicates. *p<0.05, **p<0.01, ***p<0.001. F) Representative bright-field images of intestinal organoids co-cultured with *Col6a1*-GFP^+^ and GFP^−^ IMCs at day 3, in the absence of R-Spondin 1 (Scale bar: 100 μm). G) Mean expression (z-score) of genes signatures extracted from the different intestinal mesenchymal clusters identified in Kinchen et al., [14] in *Col6a1*-GFP^+^ bulk RNA-seq samples. MF, myofibroblasts; SMC, smooth muscle cells. H) Heatmap of the top 50 differentially expressed genes in the Str2 population (Kinchen et al., [14]) and their relative expression in the GC and TC samples.

A similar cell sorting and sequencing approach was also performed for the small intestine, where two populations of GFP^+^ cells were identified, a GFP^hi^ (GIH) and a GFP^lo^ (GIL) population, each accounting for 23% and 29% of the Lin^−^ population, respectively (Figure S1A). Analysis of the genes identified in the colon showed that GIH cells were also enriched in genes related to epithelial differentiation and blood vessel function, while GFP^lo^ (GIL) cells were similar to the GFP^−^ population (Figure S1B). The presence of GFP^lo^ cells in the small intestine indicates a GFP^+^ cell population with potentially different cellular properties [22], and could represent a distinct cell subset, an intermediate state between GFP^−^ and GFP^+^ cells or the result of non-specific recombination. These results indicate that the *Col6a1^Cre^* mouse targets a broader set of mesenchymal subsets in the small intestine in relation to the colon.

Interestingly, one of the genes enriched in the GFP^−^ samples in both the small intestine and colon was *Cd34* (Figure 1B). Stzepourginski et al. [17] described a broad population of PDPN^+^CD34^+^ cells, which are located at the bottom of the crypts, express Gremlin1, Wnt2b and R-spondin, and play a role in stem cell maintenance. FACS analysis and immunohistochemistry showed that indeed the majority of GFP^+^ cells in the colon (84%) and GFP^hi^ in the small intestine (82%) were CD34 negative and further indicated a preferential subepithelial localization for GFP^+^ cells outside the bottom of the crypts (Figure 1C-D and Figure S1C-D). Consistent with this data, co-culture of either GFP^+^ or GFP^−^ colonic IMCs with intestinal organoids showed that GFP^+^ cells were less potent than GFP^−^ cells in supporting the growth of epithelial organoids in the absence or presence of R-spondin 1 (Figure 1E-F). Comparison of the gene signature of mesenchymal subsets from the recently published single-cell RNA sequencing of the mouse colon by Kinchen et al., [14] and the gene expression profile of GFP^+^ cells revealed an association with the Str2 cluster (Figure 1G-H). Importantly, this cluster appears to be the highest conserved between mice and humans [14]. These results show that the *Col6a1^Cre^* mouse targets mostly CD34^−^ mesenchymal cells outside the bottom of the colonic crypt, rendering it appropriate for the functional characterization of these cells.

### Colonic *Col6a1^Cre^*-GFP^+^ cells are CD201 positive and include PDGFRα^hi^ mesenchymal cells and perivascular cells

We next searched our sequencing data for markers that could be used to detect, isolate and study *Col6a1^Cre^*-GFP^+^ cells. We found that GFP^+^ cells express higher levels of CD201 (Procr or EPCR), which was further verified by immunohistochemistry (Figure 2A). FACS analysis indeed showed that 67% of the Lin^−^GFP^+^ population in the colon were CD201^+^(Figure 2B). It should be noted that CD201 is also expressed by endothelial cells, which are excluded as Lin^−^ in our analysis. Since not all GFP^+^ cells express CD201, we also isolated Lin^−^GFP^+^CD201^+^ and Lin^−^GFP^+^CD201^−^ cells from the colon by FACS sorting. qPCR analysis showed that GFP^+^CD201^+^, but not GFP^+^CD201^−^ cells, expressed genes related both to the regulation of blood vessel function and epithelial cell differentiation, similar to GFP^+^ cells (Figure 2C). These results suggest that the remaining *Col6a1^Cre+^*CD201^−^ cells either have another yet unknown role in the intestine or are the result of non-specific targeting by the *Col6a1^Cre^* mouse.

**Figure 2.**
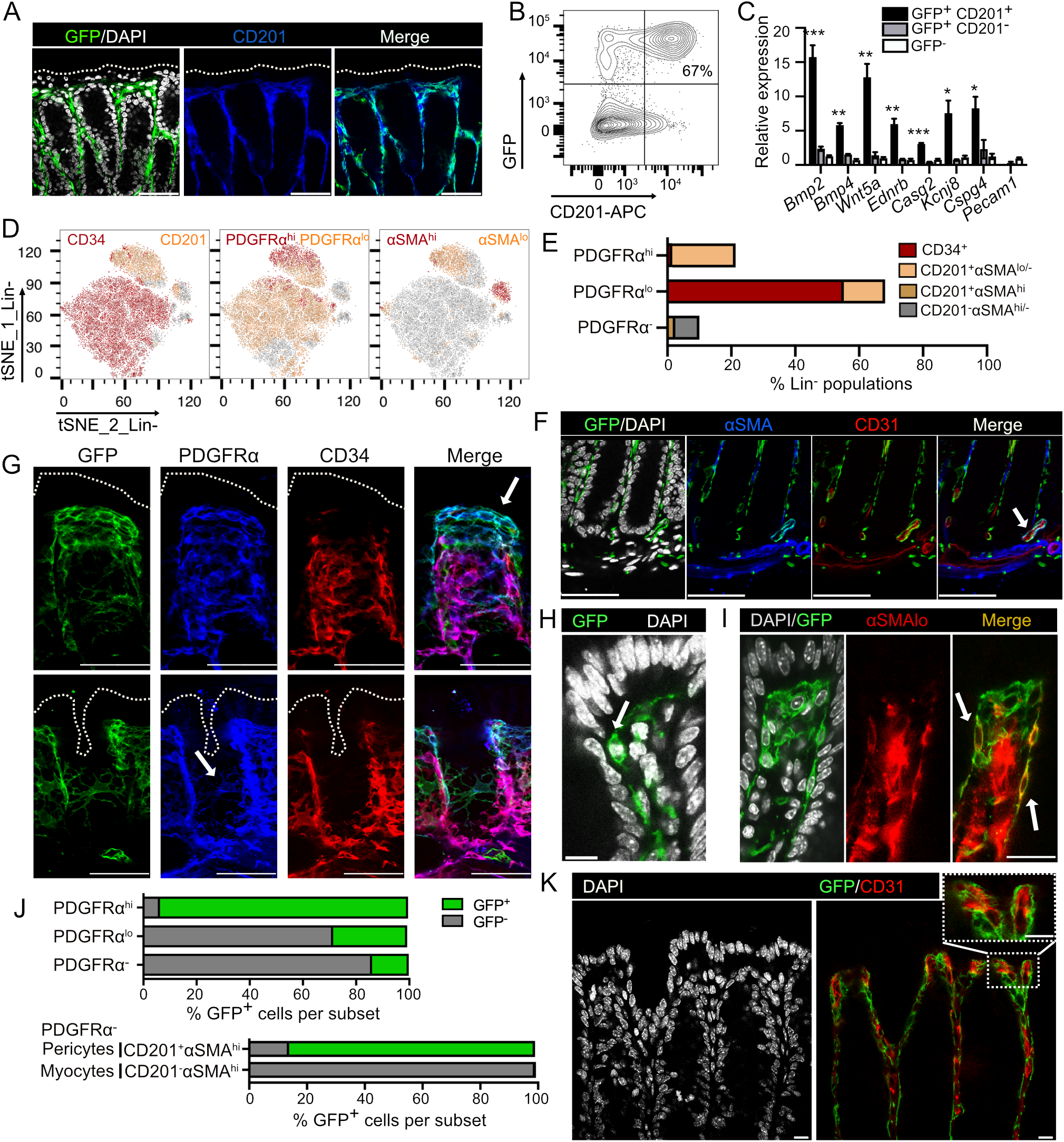
GFP^+^/CD201^+^ mesenchymal cells comprise distinct subsets in the mouse colon. A) Immunohistochemistry for CD201 in the colon of *Col6a1^mTmG^* mice (n = 4 mice, Scale bar: 50 μm). The dotted line delimitates the epithelial surface towards the lumen. B) Representative FACS analysis of CD201 expression in Lin^−^ cells in the colon of *Col6a1^mTmG^* mice (n = 10 mice). C) Gene expression analysis of selected genes in FACS-sorted GFP^+^CD201^+^, GFP^+^CD201^−^ and GFP^−^ colonic mesenchymal cells from *Col6a1^mTmG^* mice. Expression is measured in relation to the *Hprt* housekeeping gene (n = 3), *p<0.05, **p<0.01, ***p<0.001. D) tSNE plots showing the expression of CD34, CD201, PDGFRα and αSMA in Lin^−^ colonic mesenchymal cells using FACS analysis and E) quantification of CD34^+^ and CD201^+^ cells in PDGFRα^+^ subsets (n = 8 mice). F) Immunohistochemistry for αSMA and CD31 in the colon of *Col6a1^mTmG^* mice, showing their localization around blood vessels (white arrow) (Scale bar: 50 μm). The bottom of crypts is shown. G) Immunohistochemistry for PDGFRα and CD34 in the colon of *Col6a1^mTmG^* mice. Different planes are shown. White arrows indicate PDGFRα^hi^ (upper panel) and PDGFRα^lo^ (lower panel) mesenchymal cells (Scale bar: 50 μm). The dotted line delimitates the epithelial surface towards the lumen. H) Confocal imaging of GFP^+^ cells at the top of *Col6a1^mTmG^* colonic crypts (Scale bar: 10 μm). I) Immunohistochemical analysis of αSMA^lo^ cells in the colon. White arrows indicate colocalization with GFP along the crypt’s length. (Scale bar = 10 μm) (n = 4 mice). J) Quantification of *Col6a1^Cre^*-GFP^+^ cells in PDGFRα^+^ (top) and PDGFRα-(bottom) mesenchymal subsets using FACS analysis (n = 6 mice). K) Immunohistochemistry for CD31 in the colon of *Col6a1^mTmG^* mice (n = 3 mice, Scale bar: 10 μm).

Combination of the CD201 and CD34 markers was able to distinguish between the two distinct Lin^−^ mesenchymal subpopulations (Figure 2D and Figure S2A). Additional co-staining with PDGFRα and αSMA also divided PDGFRα^hi^, PDGFRα^lo^, and PDGFRα^−^ into CD201^+^ and CD34^+^ cell subsets (Figure 2D-E). PDGFRα^hi^ cells, which represent 22% of Lin^−^ mesenchymal cells, are predominantly CD201^+^αSMA^lo/-^ (93%), while PDGFRα^lo^ cells, which account for 68% of Lin-mesenchymal cells, include 81% CD34^+^ and 19% CD201^+^αSMA^lo/-^ cells (Figure 2D-E). PDGFRα^−^ cells are 10% of the Lin^−^ mesenchymal cells and include a CD201^+^αSMA^hi^ subset (24%) (Figure 2D-E and Figure S2B-D). Notably, CD34^+^ cells were mainly PDGFRα^lo/-^ and only a limited number expressed PDGFRα^hi^ and αSMA, in line with recent reports [7, 17] (Figure 2D-E).

GFP^+^PDGFRα^−^CD201^+^αSMA^hi^ cells were found around blood vessels and were most possibly pericytes (Figure 2F and Figure S2D), as also indicated by qPCR analysis (Figure S2E) and previous results [4, 23]. GFP^+^PDGFRα^hi^αSMA^lo/-^ were long thin cells in a subepithelial location found both clustered at the top of the colonic crypts and individually along the crypt’s length, corresponding to telocytes or subepithelial myofibroblast (SEMFs) (Figure 2G-I and Figure S2C, S2F) [4, 7, 13, 19]. Indeed, GFP^+^ cells expressed markers related to telocytes, including *Foxl1*, *Bmp7, Wif1* and *Wnt5a* (Figure 1B and Figure S2E). Finally, GFP^+^PDGFRα^lo^-αSMA^lo/-^ cells appeared as flat cells with extended processes that surround the colonic crypt (Figure 2G). FACS-based quantification of *Col6a1^Cre^*-GFP^+^ cells in these subsets revealed targeting of almost all the PDGFRα^hi^ cells (93.2%), and one third of the PDGFRα^lo^ stroma (27.8%). Absence of *Grem1* expression (Figure 1B) indicates that GFP^+^PDGFRα^lo^ cells do not include trophocytes. The *Col6a1Cre* mouse also targeted 13.8% of PDGFRα^−^ cells, comprising mainly CD201^+^αSMA^hi^ pericytes (86%) (Figure 2J). However, the apparent low abundancy of this population suggests that expression of genes related to blood vessel function could be also associated with the localization of *Col6a1^Cre^*-GFP*^+^* cells in close proximity to the subepithelial capillary network, indicating thus a potential dual function for colonic telocytes/SEMFs (Figure 2K). These results reveal the complexity of mesenchymal cells surrounding the colonic crypt, provide a detailed characterization of their markers and localization and the efficiency of their targeting by the *Col6a1^Cre^* mouse.

### *Col6a1^Cre^*-GFP cells are found as mesenchymal aggregates during development and orchestrate intestinal morphogenesis

To examine the specificity of *Col6a1^Cre^* mice also during embryonic organogenesis, we analysed *Col6a1^mTmG^* mice at different developmental stages. We found that GFP^+^ cells were absent until E13.5 and started to appear at E14.5-E15.5 as aggregates beneath the epithelial layer (Figure 3A). As villi became more elongated, GFP^+^ cells extended toward the bottom of the crypts (Figure 3B). GFP^+^ aggregates expressed PDGFRα, as shown by both confocal microscopy and FACS analysis, in line with previous reports [24–26] (Figure 3A-B and 3D). Notably, similar GFP^+^ PDGFRα^+^ aggregates were also detected in the developing colon (Figure 3A-B). FACS analysis showed that the majority of GFP^+^ cells corresponded to 28% and 38.3% of PDGFRα^hi^ cells at E16.5 and E18.5, respectively (Figure 3D and E). Contrary to adulthood, the *Col6α1^Cre^* mouse targeted a limited number of other mesenchymal populations, including PDGFRα^lo^ and αSMA^+^ cells, which indicates its specificity for telocytes or telocyte precursors during development (Figure 3D-Ε). It should also be noted that CD34 was not expressed at this stage in agreement with previous reports [17]. CD201 was broadly expressed, although it did include PDGFRα^hi^ cells also targeted by the *Col6α1^Cre^* mouse (Figure 3F). These results show a potential ontogenic relationship between telocytes and mesenchymal villous and colonic clusters.

**Figure 3.**
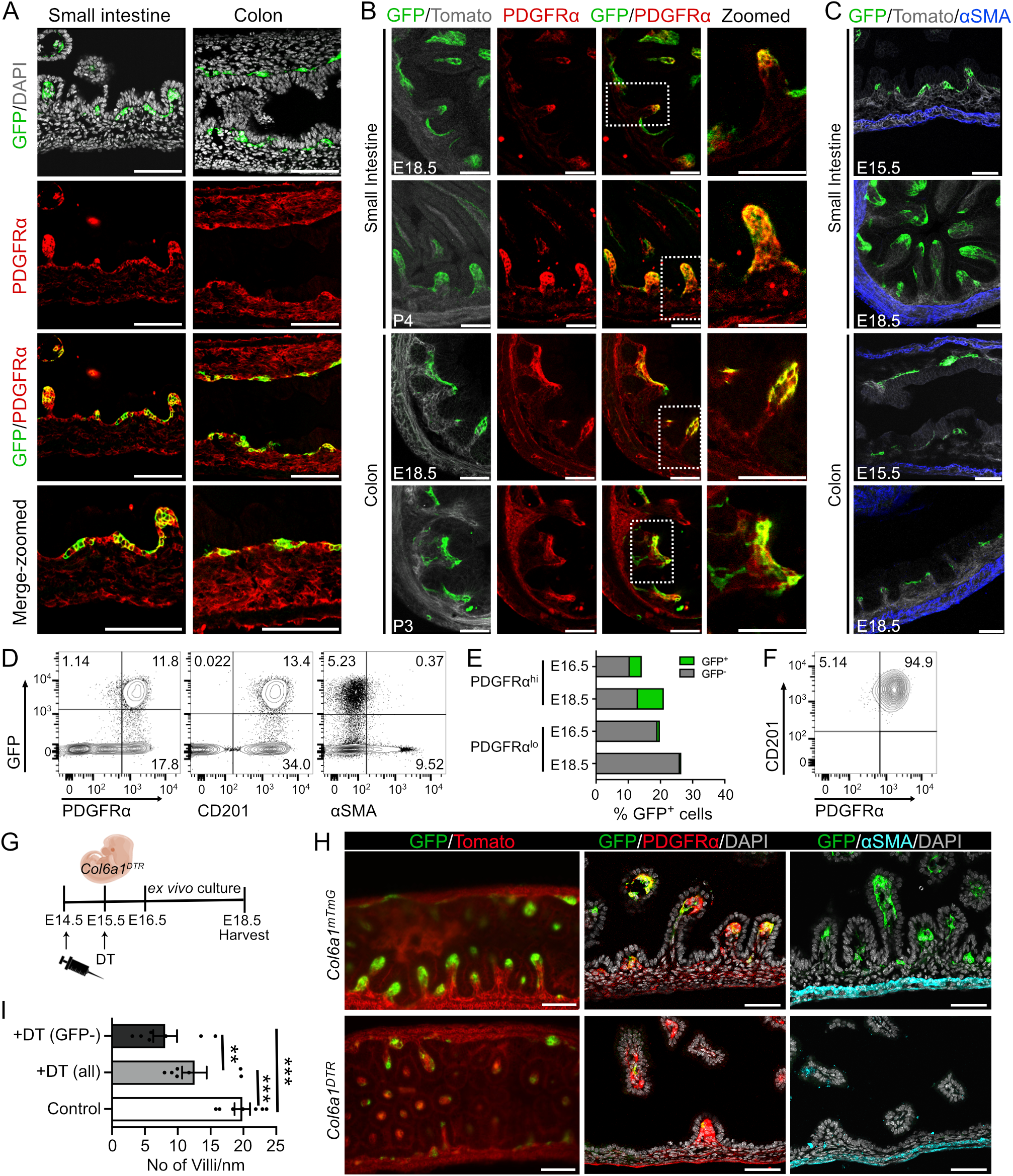
The *Col6a1^Cre^* mouse targets mesenchymal cell aggregates during development, which are necessary for intestinal morphogenesis. Confocal images showing GFP and PDGFRα expression in A) the small intestine and colon at E15.5 (Scale bar = 50 μm) and B) the small intestine and colon of *Col6a1^mTmG^* mice at the indicated developmental stages (Scale bar = 50 μm), (n = 3 mice per developmental stage). C) Confocal images showing GFP and αSMA expression in the small intestine and colon of *Col6a1^mTmG^* mice at the indicated developmental stages (Scale bar = 50 μm), (n = 3 mice per developmental stage). D) FACS analysis of *Col6a1*-GFP^+^ intestinal mesenchymal cells at E18.5. E) FACS-based quantification of GFP^+^ cells in PDGFRα^hi^ and PDGFRα^lo^ cells in E16.5 and E18.5. F) FACS analysis of GFP^+^ cells in E18.5, showing that they all are CD201^+^PDGFRα^hi^ cells (n = 3-4 mice per developmental stage in all FACS analyses). G) Schematic representation of DT administration. Pregnant females received two injections of diptheria toxin (DT) (*5*μg) at E14.5 and E15.5, which was followed by ex vivo culture of the intestine from E16.5 to E18.5. H) Lightsheet imaging (maximum projection, Scale bar = 100 μm) and confocal images showing GFP, PDGFRα and αSMA expression (Scale bar = 50 μm) in the small intestine of *Col6a1^DTR^* and control mice. I) quantification of villi/nm in the presence of DT (n = 7). All (GFP^+^ and GFP^−^) and only GFP^−^ villi are presented in DT treated mice. ***p<0.001, **p<0.01.

To define the physiological importance of these cells during intestinal embryonic development, we employed the iDTR strain [27] in combination with the *Col6a1^mTmG^* mice (*Col6a1^DTR^*). Administration of diptheria toxin (DT) in *Col6a1^DTR^* mice at Ε14.5 and E15.5 and subsequent ex vivo culture of the embryonic intestine for 48 hours resulted in depletion of *Col6a1*-GFP^+^ cells and a significant reduction in the number of developing villi (Figure 3G-I, Figure S3 and Supplementary Videos 1 and 2). Total and GFP^−^ villi were separately quantified in the *Col6a1^DTR^* mice to take into account potential inefficient or patchy deletion, while all villi appear to have GFP^+^ clusters in the control mice (Figure 3I and Supplementary Videos 1 and 2). Staining with PDGFRα and αSMA did not reveal major differences in their expression patterns after GFP^+^ cell depletion, in agreement with the level of *Col6a1^Cre^* targeting at this stage (Figure 3C-E and 3H). These results suggest that *Col6a1*-GFP^+^ telocyte precursors act as orchestrators of intestinal morphogenesis and patterning.

### Colonic *Col6a1^Cre+^* IMCs regulate homeostatic epithelial proliferation and enteroendocrine cell differentiation

To define the homeostatic roles of colonic *Col6a1^Cre+^* IMCs, we then depleted *Col6a1*-GFP^+^ cells in 4-6-month-old *Col6a1^DTR^* mice. Due to increased lethality upon systemic DT injection, we performed local intrarectal DT administration, as shown in Fig. 4A. Efficient, albeit not complete, depletion of GFP^+^ cells (75 % reduction) at the last 3-4 cm of the colon was verified by confocal imaging and FACS analysis (Fig. 4B-C). Further analysis of the remaining GFP^+^ cells showed that although all subtypes were depleted, there was a preferential loss of PDGFRα^hi^ cells and a proportional increase in PDGFRα^lo^ cells (Fig. 4D). Accordingly, and consistent with our previous data (Fig. 2), PDGFRα^hi^CD201^+^ telocytes/SEMFs were reduced by 90%, while PDGFRα^lo^ stromal cells were proportionally increased by 11%, and PDGFRα^−^ cells remained largely unaltered (Fig. 4E-F). These results indicate that PDGFRα^hi^ telocytes/SEMFs is the mesenchymal subpopulation most efficiently depleted using our approach. Histopathological examination of *Col6a1^DTR^* mice showed that the intestinal structure was normal following DT administration (Figure 4G). Immunohistochemistry and/or qPCR analysis for the quantification of specific intestinal epithelial subpopulations showed a reduction in enteroendocrine cell differentiation upon GFP^+^ cell depletion, while stem cell maintenance, as well as the differentiation of Tuft and Goblet cells was not affected (Figure 4H-I and Figure S4). Quantification of telocyte markers in these tissue samples showed a reduction in *Bmp7* and *Wnt5α* expression levels, while the expression of most other Bmps and *Foxl1* was not altered (Figure 4J). Notably, Bmp7, which forms heterodimers with Bmp2 and Bmp4 was previously shown to be one of the factors that is exclusively expressed by PDGFRα^+^ telocytes at the crypt-villous boundary of the small intestine [7]. In addition, we also detected a defect in the distribution of BrdU^+^ proliferating epithelial cells along the crypt axis, characterized by an increase toward the top of the crypt, which could be associated with the role of BMPs in the regulation of stem cells [28] (Figure 4K-L). These results show that colonic *Col6a1^Cre+^* IMCs have distinct pathophysiological roles in epithelial cell differentiation and proliferation, although they are largely dispensable for normal tissue architecture and function.

**Figure 4.**
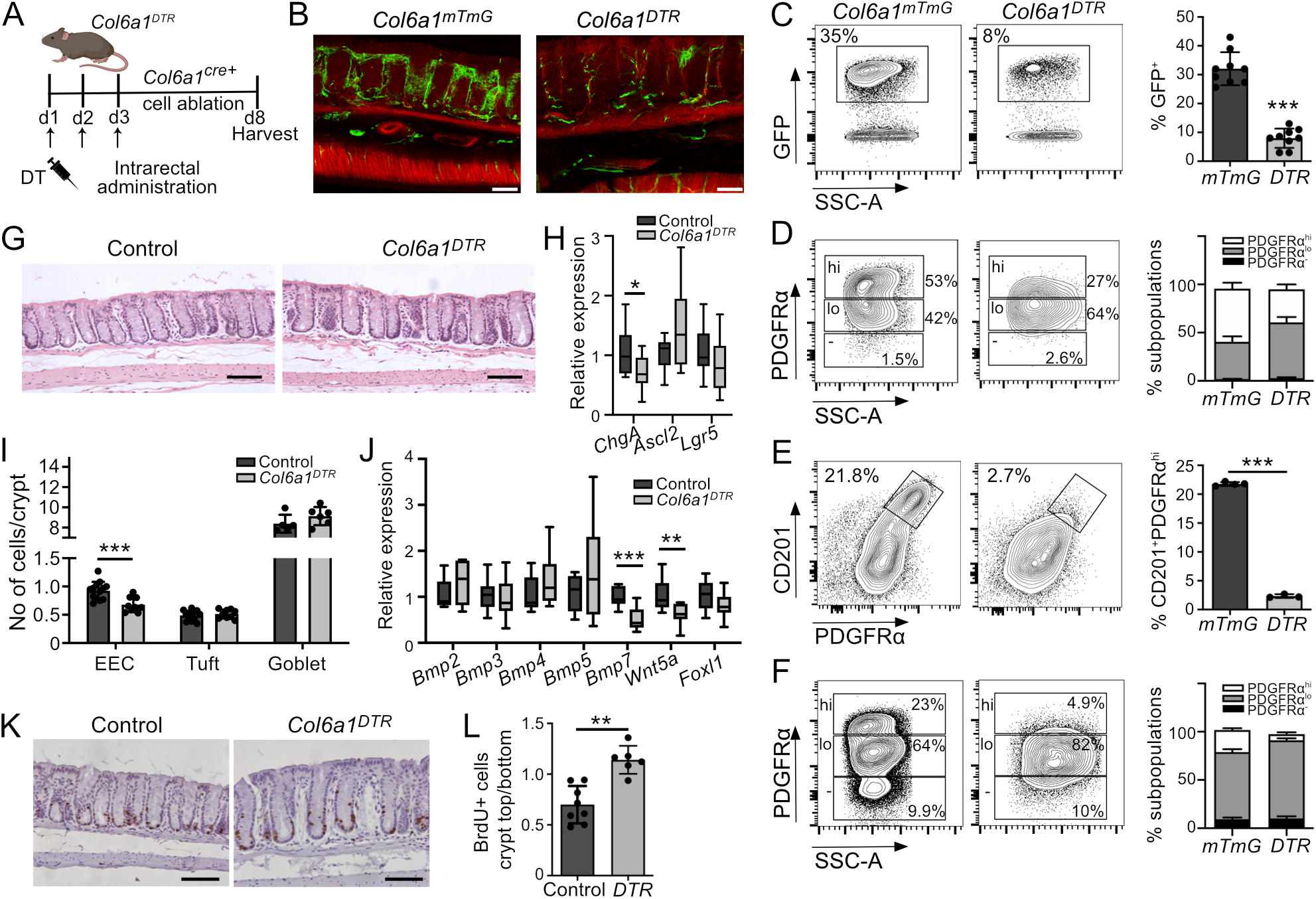
*Col6a1^Cre+^* cell depletion leads to deregulated epithelial cell differentiation and proliferation during homeostasis. A) Schematic representation of DT administration in homeostasis. *Col6a1^DTR^* and control (*Col6a1^mTmG^*, *iDTR^f/f^*^/^) mice received 3 daily intrarectal administrations of DT (*20ng/g* body weight) and mice were sacrificed after 5 days. B) Confocal images of GFP expression in *Col6a1^mTmG^* and *Col6a1^DTR^* mice (Scale bar = 50 μm). C) FACS analysis and quantification of GFP^+^ cells in the colon of *Col6a1^mTmG^* and *Col6a1^DTR^* mice after DT administration (n = 9). D) FACS analysis and quantification of PDGFRα^+^ subsets in GFP^+^ cells in *Col6a1^mTmG^* and *Col6a1^DTR^* mice (n = 3-5). E) FACS analysis and quantification of CD201^+^PDGFRα^hi^ cells in *Col6a1^mTmG^* and *Col6a1^DTR^* mice (n = 3-4 mice). F) FACS analysis of PDGFRα^+^ subsets in *Col6a1^mTmG^* and *Col6a1^DTR^* mice (n = 4-10). G) H&E staining of *Col6a1^DTR^* and control (*Col6a1^mTmG^*, *iDTR^f/f^*^/^) mice (Scale bar: 100 μm). H) Expression analysis of the indicated genes in colon samples from *Col6a1^DTR^* and control (*Col6a1^mTmG^*, *iDTR^f/f^*^/^) mice. Expression is measured in relation to the *B2m* housekeeping gene (n = 9-14). I) Immunohistochemical-based quantification of differentiated epithelial cell types per crypt in *Col6a1^DTR^* and control (*Col6a1^mTmG^*, *iDTR^f/f^*^/^) mice (n = 5-13). J) Expression analysis of the indicated genes in colon samples from *Col6a1^DTR^* and control (*Col6a1^mTmG^*, *iDTR^f/f^*^/^) mice. Expression is measured in relation to the *B2m* housekeeping gene (n = 5-15). K) Representative BrdU staining and L) quantification of the ratio of BrdU^+^ cells in the top/bottom of the colonic crypts of *Col6a1^DTR^* and control (*Col6a1^mTmG^*, *iDTR^f/f^*^/^) mice (Scale bar: 100 μm), (n = 6-7). *p<0.05, **p<0.01, ***p<0.001.

### Loss of *Col6a1^Cre+^* IMCs is followed by CD34^+^ mesenchymal cell plasticity

The similar expression levels of most BMPs in the *Col6a1^DTR^* mice and its normal tissue architecture indicated that other mesenchymal cell populations could mediate some of the functions of *Col6a1^Cre+^* IMCs under these conditions. Indeed, we found that colonic crypt tops in the *Col6a1^DTR^* mice were populated by GFP^−^ PDGFRα^+^CD34^+^ cells, which however remained PDGFRα^lo^ (Figure 5A-B). CD34^+^ cells in this area were also Ki67^+^, indicating that these cells could proliferate and occupy the space, where *Col6a1^Cre+^* telocytes/SEMFs were previously located (Figure 5C-D). They also expressed αSMA, a marker commonly upregulated in response to fibroblast activation (Figure 5E-G). Most importantly, they showed increased expression of genes associated with telocyte functions, including *Bmp7*, *Wnt5a* and *Foxl1* (Figure 5H). These results show that following *Col6a1^Cre+^* IMC depletion, CD34^+^ cells become activated, they proliferate and occupy the space at the top of the colonic crypts, partly compensating for the loss of telocytes, providing thus evidence for mesenchymal plasticity in the intestine.

**Figure 5.**
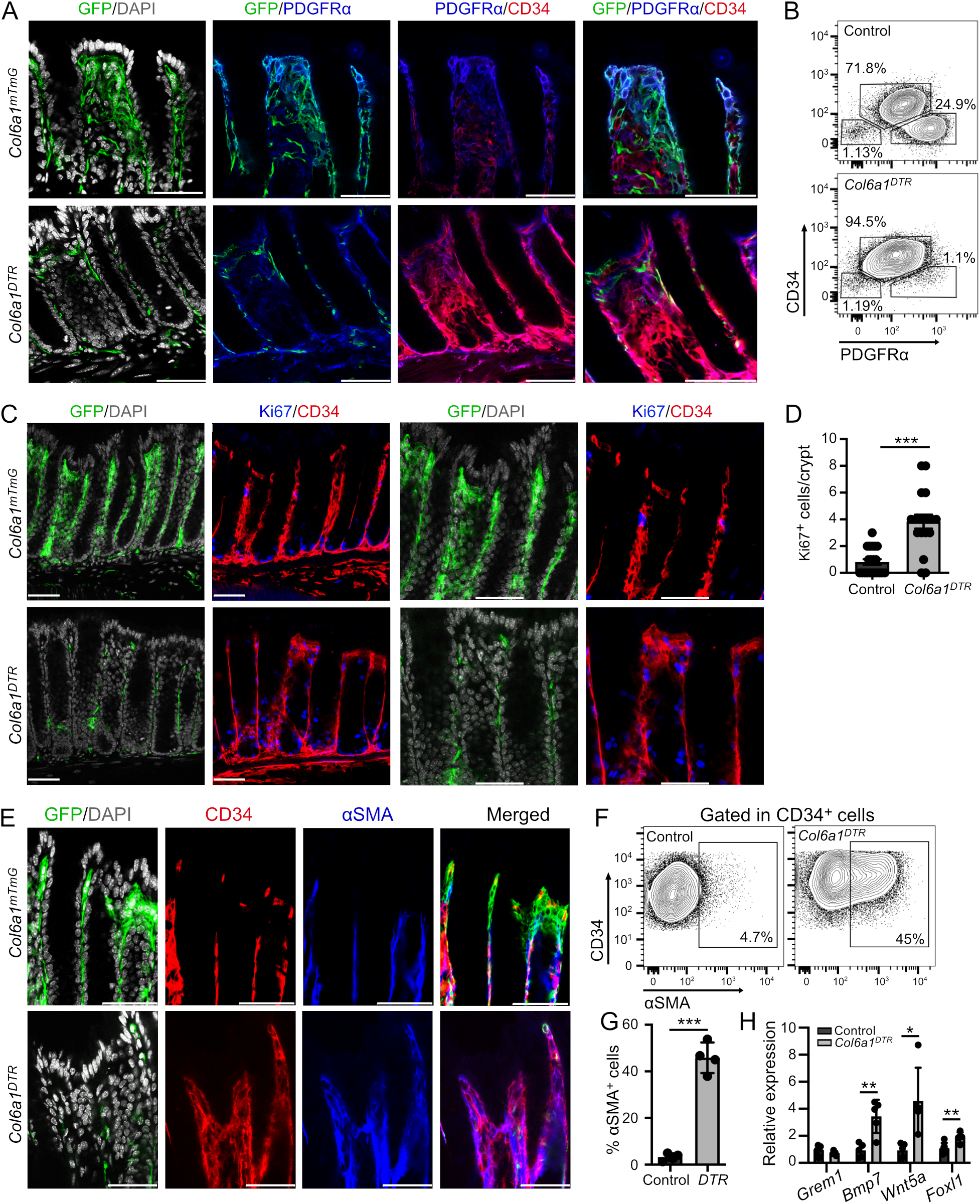
Loss of *Col6a1^Cre+^* IMCs is followed by CD34^+^ mesenchymal cell plasticity. A) Immunohistochemical and B) FACS analysis of CD34 and PDGFRα expression in the colon of *Col6a1^DTR^* and *Col6a1^mTmG^* mice (Scale bar = 50 μm) (n = 6 mice). A zoomed-in view of the crypt top is shown in the merged image. C) Immunohistochemical analysis and D) quantification of Ki67^+^ mesenchymal cells in the colon of *Col6a1^DTR^* and *Col6a1^mTmG^* mice (Scale bar = 50 μm) (n = 5 mice). Both broad and zoomed-in views of the crypt top are shown. E) Immunohistochemical analysis of αSMA expression in the colon of *Col6a1^DTR^* and control mice (the top of the crypt is shown) (n = 5). F) FACS analysis and G) quantification of αSMA^+^ cells in the CD34^+^ subset (n = 4 mice). H) Gene expression analysis of isolated CD34^+^ cells from *Col6a1^DTR^* and control mice (n = 5). *p<0.05, **p<0.01, ***p<0.001.

### Topological and functional plasticity of mesenchymal cells during intestinal regeneration

To further explore the functions of *Col6a1*-GFP^+^ cells in non-homeostatic conditions, we subjected *Col6a1^mTmG^* mice to the DSS model of acute colitis and isolated Lin-GFP^+^ (GDS) and Lin-Tomato^+^ (TDS) cells by FACS sorting, as previously described (Figure 6A). Comparisons between GFP^+^ and GFP^−^ IMCs and the unsorted population (UDS), as well as cells from untreated mice and subsequent Gene Ontology analysis revealed that both GDS and TDS cells were enriched for functions related to inflammatory/ immune responses (Figure 6B). GDS and TDS samples also showed enrichment in genes related to epithelial proliferation/differentiation and blood vessel function, similar to the homeostatic situation, suggesting that these cells, although activated, retain their homeostatic properties and marker expression (Figure 6B and Figure S5A). FACS analysis of mesenchymal populations described in Figure 2 verified their presence also during acute colitis, although CD201^+^PDGFRα^−^αSMA^hi^ were increased in accordance with fibroblast activation (Figure S5B-C). Confocal imaging further revealed that in sites of ulceration GFP^+^ cells were located in the upper part of the damaged area, indicating that they also retained their topology (Figure 6C). This was further verified by comparisons with the recently published single cell RNA-seq data by Kinchen et al., which confirmed that GDS cells corresponded mostly to the Str2 population [14] (Figure 6D).

**Figure 6.**
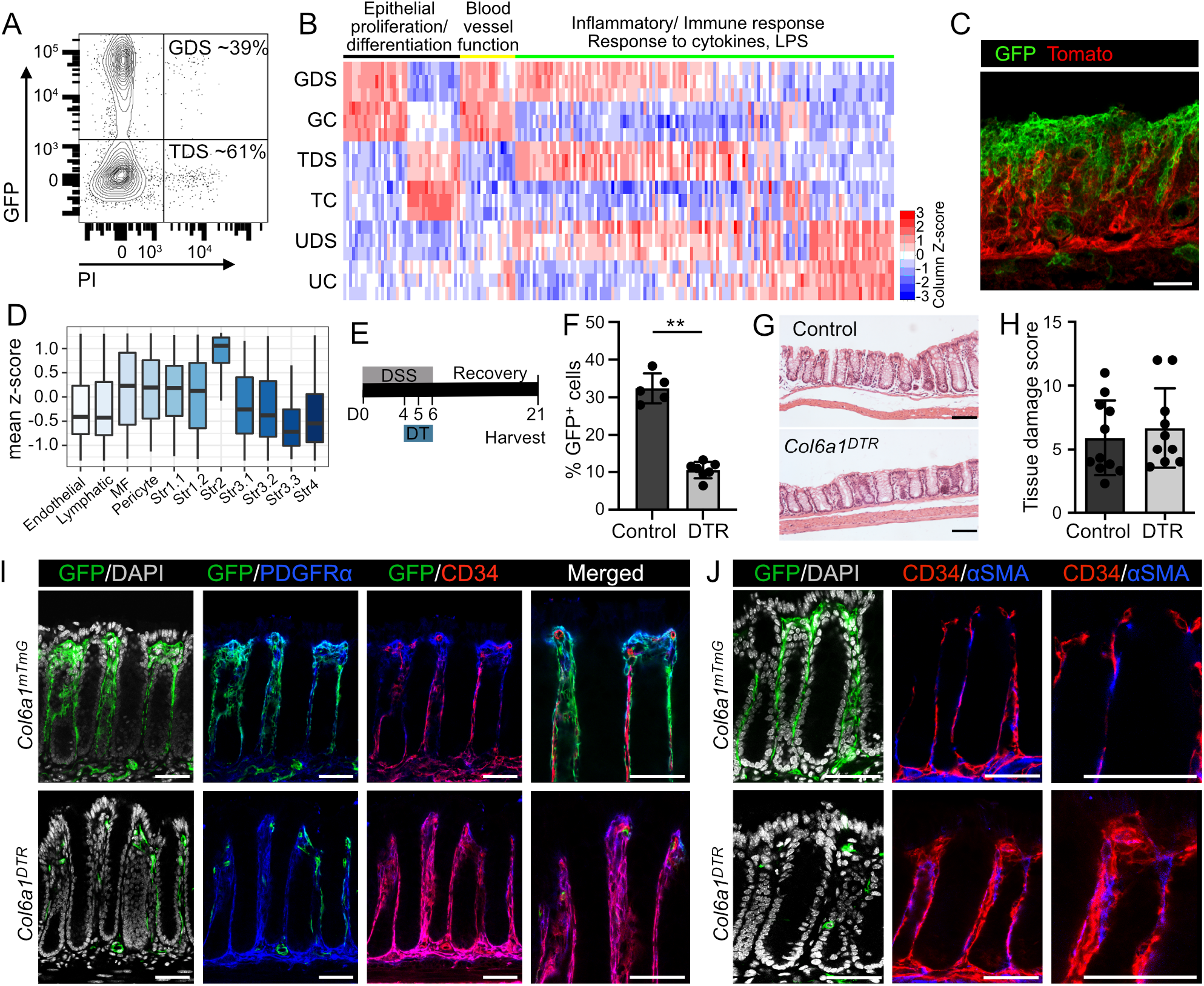
*Col6a1^Cre+^* cells retain their properties and topology during colitis but are dispensable for tissue regeneration. A) FACS sorting strategy for the isolation of *Col6a1^Cre^*-GFP^+^ (GDS) and *Col6a1^Cre^*-GFP^−^ (TDS) mesenchymal cells from the colon at the end of the acute DSS protocol. 3 samples from 4-5 mice each were subsequently analyzed. B) Heatmap of differentially expressed genes in GDS vs TDS and UDS, as well as the respective untreated samples, corresponding to GO terms related to epithelial proliferation/differentiation, blood vessel regulation and inflammatory response. Log2 transformed normalized read counts of genes are shown. Read counts are scaled per column, red denotes high expression and blue low expression values. C) Confocal images of GFP expression in *Col6a1^mTmG^* mice at day 8 of the acute DSS protocol, Scale bar = 50 μm. D) Mean expression (z-score) of genes signatures extracted from the different intestinal mesenchymal clusters identified in Kinchen et al., (14) during DSS colitis in *Col6a1*-GFP^+^ bulk RNA-seq samples. MF, myofibroblasts. E) Schematic representation of DT administration during acute colitis and regeneration. Mice received 2.5% DSS for 5 days, followed by regular water for 16 days. 100μl DT (*20ng/g* body weight) was administered intrarectally at days 4, 5 and 6 of the regime. F) Quantification of GFP^+^ cells in the colon of control and *Col6a1^DTR^* mice after DT administration (n = 5-7 mice), G) H&E staining and H) histopathological score of *Col6a1^DTR^* and control (*Col6a1^mTmG^*, *iDTR^f/f^*^/^) mice at the end of the protocol (Scale bar = 100 μm) (n = 10-11 mice). Immunohistochemical analysis of I) CD34 and PDGFRα expression and J) CD34 and αSMA expression in the colon of *Col6a1^DTR^* and control mice (Scale bar = 50 μm) (n = 4 mice).

Depletion of GFP^+^ cells during DSS administration and evaluation of tissue morphology 21 days after the initiation of the protocol showed similar histopathological score to untreated mice, indicating normal regeneration of the intestine (Figure 6E-H). Similar to homeostasis, GFP^−^PDGFRα^+^CD34^+^ cells were localized at the crypt tops, indicating the plasticity of CD34^+^ cells and their ability to support the normal re-epithelization of the colon (Figure 6I). Notably, at this late time-point, CD34^+^ cells were not αSMA^+^, indicating their potential reversible activation following *Col6a1Cre^+^* cell depletion (Fig. 6J). Therefore, these results further support the plasticity of intestinal mesenchymal cells and suggest that reciprocal signals between the epithelium and the underlying mesenchyme are more or equally important to specialized cell types for intestinal regeneration.

## Discussion

The importance of the mesenchymal stroma in the maintenance of intestinal structure and function, as well as in the response to injury is now well-established [3, 4]. Opposing signaling gradients along the colonic crypt length and the crypt/villous axis in the small intestine, including predominantly Wnts, BMPs, and their inhibitors, are considered crucial for the maintenance of this morphology and the presence of specialized mesenchymal cell types could explain this phenomenon [8]. Indeed, several studies have described specific cell subsets that act as regulators of the stem cell niche [6, 7, 13, 15–18]. Accordingly, although less studied, telocytes and specific fibroblast subsets identified by sc-RNA sequencing analyses have been shown to express BMPs and regulators of epithelial differentiation [6, 7, 14, 19, 29].

In this study, we focused on characterizing the identities, markers, origins and functional significance of IMCs outside the colonic stem cell niche in intestinal development, homeostasis and tissue damage/regeneration using the *Col6a1^Cre^* transgenic mouse [30]. To this end, we have performed transcriptomic, FACS and imaging analysis of the cells targeted by the *Col6a1^Cre^* mouse strain. We also identified a novel extracellular marker, CD201, and in combination with PDGFRα and αSMA we were able to show that the *Col6a1^Cre^* mice target almost all CD201^+^PDGFRα^hi^αSMA^lo/-^ telocytes/SEMFs, around one third of the PDGFRα^lo^ stroma, and specifically CD201^+^PDGFRα^lo^ cells, as well as perivascular cells that are CD201^+^αSMA^hi^. Among them, PDGFRα^hi^ telocytes/SEMFs are the only subpopulation targeted to its entirety, and it was shown to be present along the colonic crypts, concentrating at their tops, in a similar fashion to the small intestine [7]. In addition, they were in close proximity to the subepithelial capillary network, indicating a potential dual role of telocytes/SEMFs in both epithelial and endothelial cell function.

Little is known about the developmental origins of the different mesenchymal subsets, including those identified through multiple single-cell analyses. Our results showed that *Col6a1^Cre+^* mice also target mesenchymal clusters during development both in the small intestine and colon, indicating an ontogenic relationship between them and telocytes. These clusters have been previously shown to emerge in the murine small intestine in waves after E13.5, they are PDGFRα^+^, respond to Hedgehog signaling and express BMPs to regulate intestinal morphogenesis [10, 24–26]. Similar mechanisms play an important role also in chick intestinal development, where smooth muscle differentiation drives villi formation through forces that generate localized pockets of high Shh, crucial for the expression of mesenchymal cluster genes, such as PDGFRα and BMP4 [31, 32]. Deletion of either PDGFα or PDGFRα in mice was shown to result in abnormal villi development due to reduced mesenchymal proliferation [26]. Accordingly, inhibition of Hedgehog signaling led to lack of cluster formation [24]. However, PDGFRα is broadly expressed in the intestinal mesenchyme also during development, including both high- and low-expressing cells, as we also show [26, 33]. Similarly, the Ptc1 receptor and the downstream regulator Gli are not exclusive to mesenchymal aggregates during development [24]. FoxL1 deletion also leads to a delay in vilification, but it is expressed earlier at E12.5 and could target additional populations [34]. In contrast, during development, the *Col6α1^Cre^* mouse targets exclusively a fraction of PDGFRα^hi^ cells located at the top of villi and colonic crypts. Therefore, our functional analysis through cell depletion experiments shows the significance of mesenchymal clusters and possibly telocytes precursors in intestinal villification and patterning. Notably, a similar PDGFRA^hi^ population was also recently described in the human developing intestine [35].

Interestingly, depletion of this population during homeostasis did not affect the architecture and morphology of the colon. As we show, this could be explained by the plasticity of CD34^+^ cells that are able to proliferate and occupy the area, where *Col6a1^Cre+^*/CD201^+^PDGFRα^hi^ cells were previously found. However, several aspects of intestinal homeostasis were compromised, including the expression levels of *Bmp7* and *Wnt5a*, the normal differentiation of enteroendocrine cells and the distribution of proliferating cells along the crypt axis. Both the function of enteroendocrine cells and the stemness of Lgr5^+^ cells have been previously shown to be modulated by BMP signaling [9, 28], including Bmp7, which forms heterodimers with Bmp2 and Bmp4 [36]. The exact CD34^+^ cell subset that displays such plasticity is yet not known; however, Gremlin-1^+^ cells have been previously shown to have stem cell potential in the intestine [37]. The phenotype of *Col6a1^Cre+^*/CD201^+^ IMC depletion is in contrast to FoxL1^+^ telocyte deletion using a similar methodology. However, the *Foxl1^Cre^* mouse has been shown to also target pericryptal telocytes that express Gremlin-1 [6, 13]. Although a small population, Gremlin-1^+^ cells are crucial for intestinal homeostasis and structure, as recently shown [7]. Therefore, our results, delineate the functional importance of *Col6a1^Cre+^* IMCs at the top of the colonic crypts, illustrate how they can affect epithelial cell differentiation and proliferation and provide evidence for mesenchymal plasticity towards tissue homeostasis.

Similar to homeostasis, *Col6a1^Cre+^* cell depletion during acute DSS colitis did not affect inflammation and regeneration of the intestine, despite their inflammatory gene signature and specific topology at the top of the ulcerated tissue and thus their proximity to the regenerating epithelium. As we show, this is also associated with the plasticity of CD34^+^ mesenchymal cells, which can replace *Col6a1^Cre+^* IMCs during the re-epithelization of the intestine, although the precise identities of these cells remain unknown. These results further suggest that reciprocal communication of the mesenchyme with the regenerating epithelium is crucial to orchestrate tissue regeneration, although the specific molecular pathways are not yet clear.

In conclusion, we have described the properties and identities of *Col6a1^Cre+^* colonic IMCs, including telocytes/SEMFs and perivascular cells, defined their ontogenic relationship with mesenchymal aggregates and identified their role as orchestrators of intestinal morphogenesis and regulators of epithelial homeostasis. We have further introduced the concept of mesenchymal plasticity both during homeostasis and tissue repair. In the future, it would be interesting to characterize the identities and role of these cells also in other intestinal disorders, including cancer, and define the molecular mechanism driving mesenchymal plasticity in homeostasis and disease.

## Materials and Methods

### Mice and Study Approval

*Col6a1^Cre^* mice were described before [30]. *Rosa26^mT/mG^* and *Rosa26^iDTR^* mice were purchased from the Jackson Laboratory [22, 27]. All mice were maintained under specific pathogen free conditions in the Animal House of the Biomedical Sciences Research Center “Alexander Fleming”. All studies were performed according to all current European and national legislation and approved by the Institutional Committee of Protocol Evaluation in conjunction with the Veterinary Service Management of the Hellenic Republic Prefecture of Attika under the permissions 5759/15, 8443/17, 993/18, 448195/19.

### DSS colitis induction

DSS-induced colitis was performed as previously described [38]. Briefly, 6-10 month old mice received 2.5% DSS in their drinking water, followed by 1-14 days of regular water. Colitis induction was monitored by measuring weight loss.

### Diptheria Toxin experiments

*Col6a1^DTR^*, control iDTR and *Col6a1^Cre^* mice were subjected to intrarectal administration of 100μl diphtheria toxin (Sigma-Aldrich) dissolved in 0.9% sodium chloride at 20 ng/g body weight. Pregnant mice were injected i.p. for two days (E14.5 and E15.5) with 5μg DT.

### Embryo manipulation

Timed pregnancies were set up by checking vaginal plugs to obtain E13.5, E14.5, E16.5 and E18.5 embryos. Pregnant females were sacrificed by cervical dislocation on the specific post coitum day and embryos were dissected in ice-cold PBS. Embryos were fixed overnight in 4% PFA/PBS, immersed in serial solutions of 15% and 30% sucrose/PBS and embedded in OCT for cryosection preparations.

### Isolation and culture of IMCs

Isolation and culture of IMCs was performed as previously described [39]. Briefly, the colon or small intestine was removed and digested as described above. The cell pellet was resuspended in culture medium, consisting of DMEM (Biochrom), 10% FBS (Biochrom), 100 U/mL penicillin/100 mg/mL streptomycin (Gibco), 2 mM L-Glutamine (Gibco), 1 μg/ml amphotericin B (Sigma) and 1% non-essential amino acids (Gibco) and plated in cell culture flasks. The medium was changed after 3-24 hours and cells were used after 2-4 days.

### Lightsheet Microscopy

For Lightsheet microscopy, the intestine of the embryos was isolated and fixed in 4% PFA/PBS overnight. Tissue clearing was achieved using the Scale A2 clearing solution for 2 weeks [40]. The Lightsheet Z.1 from ZEISS, equipped with sample chamber and Clr Plan-Apochromat 20x/1.0, Corr nd=1.38 lens was used for experiments with tissue cleared by Scale medium, which has a refractive index of n=1.38. Quantification of villi was performed using the Imaris Software.

### Immunohistochemistry

For confocal microscopy and immunohistochemistry, mice were perfused with 4% PFA prior to the resection of the colon or small intestine. The tissue was then incubated in 4% PFA/PBS overnight and either immersed in serial solutions of 15% and 30% sucrose/PBS and embedded in OCT (VWR Chemicals) for cryosection preparations or in 2% agarose for sectioning with a vibratome (Leica). Sections were subsequently blocked using 1%BSA in TBS containing 0.05% Tween 20 (Sigma) (TTBS) and stained with antibodies listed in Table 1. Staining for CD201 was performed in unfixed tissue, embedded in 4% agarose in CO_2_ independent medium (ThermoFisher) and sectioned with vibratome. Sections were blocked in 5% FBS in CO2 independent medium and stained with anti-CD201 antibody. 2hr fixation in 4% PFA/PBS followed prior to the addition of secondary antibody. Mounting medium containing DAPI (Sigma-Aldrich) was used to stain the nuclei. Images were acquired with a Leica TCS SP8X White Light Laser confocal system.

**Table 1.**
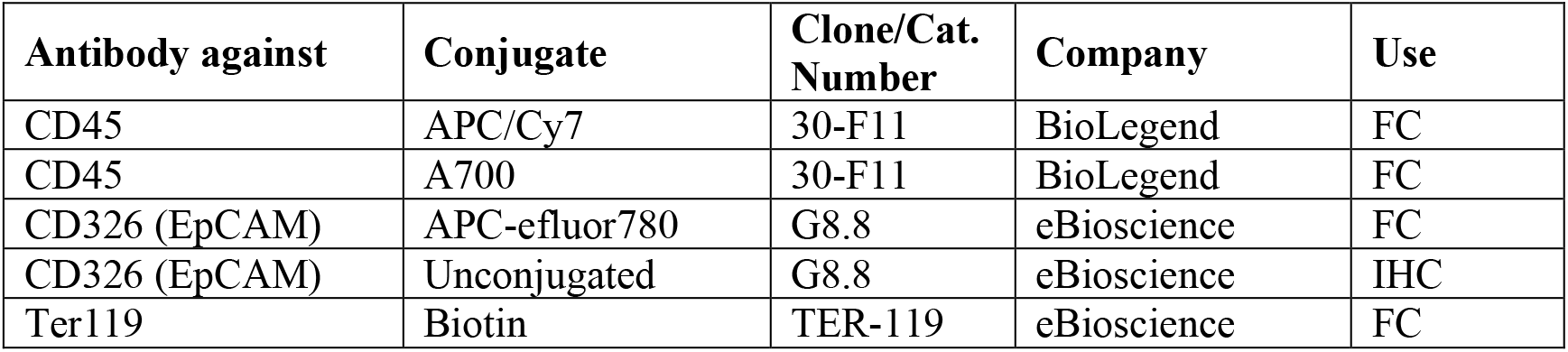
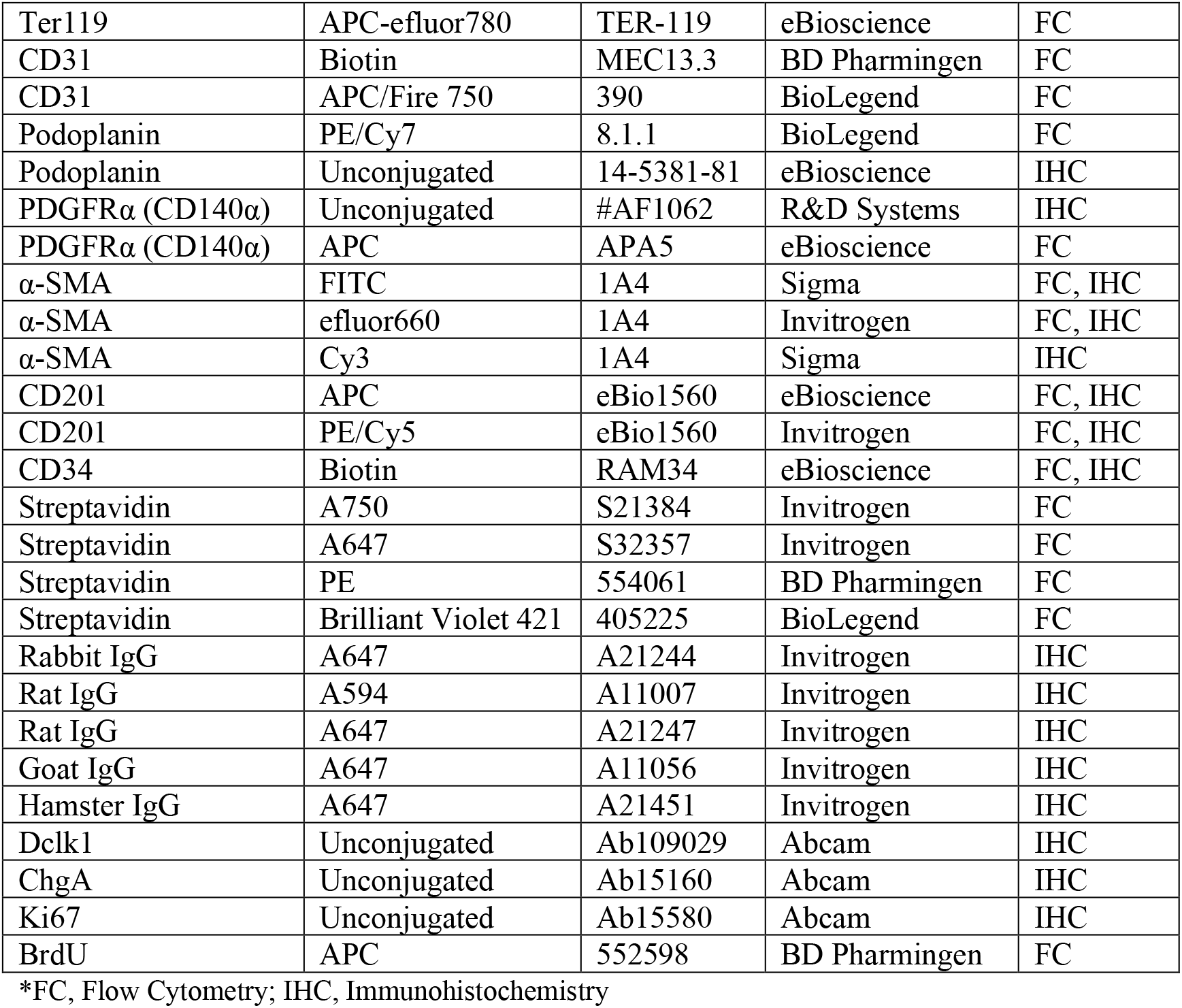
Antibodies used in flow cytometry and immunohistochemistry.

For histopathology, colon tissues were fixed in 10% formalin and embedded in paraffin. FFPE sections were stained with hematoxylin (Sigma-Aldrich) and eosin (Sigma-Aldrich) and colitis score assessment was performed as previously described [41]. Stainings for epithelial cell differentiation markers were performed in FFPE section using the antibodies listed in Table 1. Signal detection and development were performed using Vectastain ABC-HRP Kit and ImmPACT DAB kit (Vector laboratories). Quantification of proliferating cells was performed in mice that were injected i.p. with 100 mg/kg BrdU (Sigma-Aldrich) 2h prior to sacrificing them, using the BrdU detection kit (BD), according to the manufacturer’s instructions. The number of BrdU^+^ cells was quantified in at least 30 intact, well-oriented crypts per mouse.

### FACS analysis and sorting

Intestinal tissue preparations were prepared as previously described [21]. Briefly, colon or small intestine was removed, flushed with HBSS (Gibco), containing antibiotic-antimycotic solution (Gibco) and cut into pieces. Intestinal pieces were incubated with HBSS, containing 5mM EDTA, DTT and Penicillin/Streptomycin (Gibco) for 30 min, at 37°C, to remove epithelial cells. After vigorous shaking, the remaining pieces were digested using 300 U/ml Collagenase XI (Sigma-Aldrich), 1 mg/ml Dispase II (Roche) and 100 U/ml Dnase I (Sigma-Aldrich) for 40-60 minutes at 37°C. For embryos, the intestine of the embryos was isolated, cut into pieces and digested using 100 μg/ml Collagenase P (Roche), 800 μg/ml Dispase II (Roche), 200 μg/ml Dnase I (Sigma-Aldrich) for 20 min at 37°C. The cell suspension was passed through a 70μm strainer, centrifuged and resuspended in FACS buffer (PBS with 2% FBS). For stainings, 1-2 million cells/ 100μl were incubated with the antibodies shown in Table 1. For intracellular stainings, cells were fixed and permeabilized using the Fixation and Permeabilization Buffer Set (eBioscience), according to manufacturer’s instructions. Propidium Iodide (Sigma) or the Zombie-NIR Fixable Viability Kit (Biolegend) was used for live-dead cell discrimination. Samples were analyzed using the FACSCanto II flow cytometer (BD) or the FACSAria III cell sorter (BD) and the FACSDiva (BD) or FlowJo software (FlowJo, LLC).

### Crypt isolation and co-culture with IMCs

Intestinal crypts were isolated as described previously [42]. Briefly, the small intestine was flashed with cold PBS (Gibco), opened longitudinally and villi were scraped off using a coverslip. Then, it was cut into 5mm pieces and washed extensively until the supernatant was clear. Ice-cold crypt isolation buffer (2 mM EDTA in PBS) was added to the fragments and stirred for 1 hour at 4°C. Fragments were allowed to settle down, the supernatant was removed and ice-cold 2 mM EDTA/PBS was added followed by pipetting up and down. Released crypts were passed through a 70-μm-cell strainer and the procedure was repeated until most of crypts were released. Crypt fractions were centrifuged at 300*g* for 5 minutes and resuspended with ice-cold basal culture medium (Advanced DMEM/F12 (Gibco) supplemented with 2 mM GlutaMax (Gibco), 10 mM HEPES (Gibco) and 100 U/mL penicillin/ 100 mg/mL streptomycin (Gibco). Crypts were centrifuged again at 200*g* for 5 minutes, resuspended in warm basal culture medium and counted. Crypts were mixed with Col6a1cre-GFP^+^ and GFP^−^ IMCs sorted by FACS at passage 1 and subsequently resuspended in Matrigel (BD Biosciences) at 250 crypts/50.000 IMCs/30 μl in 48-well plates. After Matrigel polymerization, culture medium was added in the wells, consisting of DMEM/F12 medium (Gibco), Glutamax (Gibco), Penicillin/Streptomycin, N2 supplement (Life Technologies, 1×), B27 supplement (Life Technologies, 1×), and 1 mM N-acetylcysteine (Sigma-Aldrich), 50 ng/ml EGF (Life Technologies), 100 ng/ml Noggin (PeproTech) and Rspo1 (PeproTech) where indicated. Images were acquired with the Zeiss Axio Observer Z1 microscope. Organoid measurements were performed using the ImageJ/Fiji software.

### RNA isolation and qRT-PCR

RNA was isolated using the RNeasy mini kit or the RNeasy micro kit (Qiagen), depending on the number of cells, according to the manufacturer’s instruction. 100ng-1μg of RNA was used to generate cDNA using the MMLV reverse transcriptase by Promega and oligo-dT primers (Promega), according to the manufacturer’s instructions. For qRT-PCR, the SYBR Green PCR Master Mix (Invitrogen) was used according to the manufacturer’s instructions. Forward and reverse primers were added at a concentration of 0.2 pmol/ml in a final volume of 20 μl and qRT-PCR was performed on a CFX96 Touch™ Real-Time PCR Detection System (Bio-Rad). The primer list can be found in Table 2.

**Table 2.**
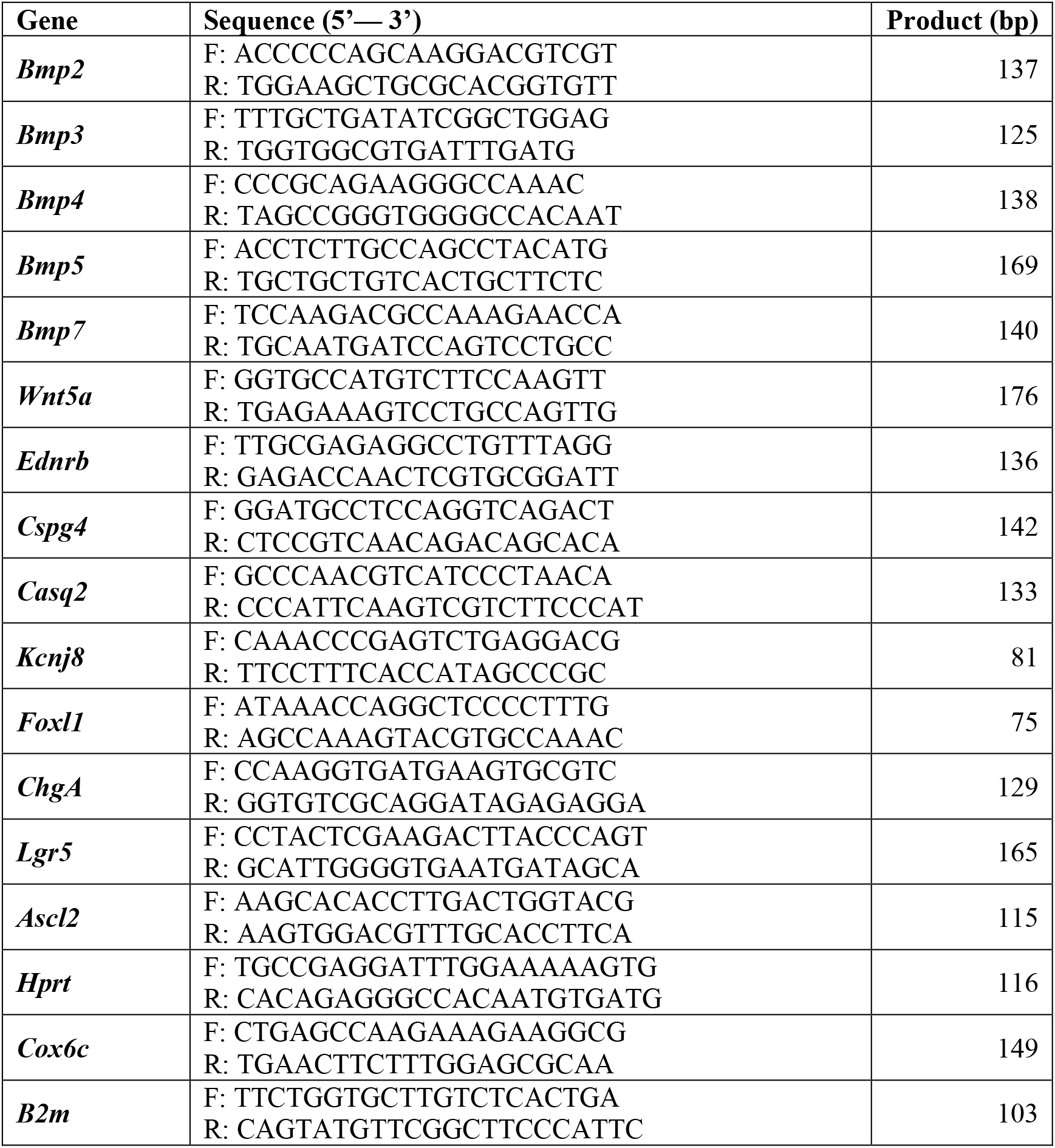
List of primers used for real-time PCR.

### 3’ RNAseq sequencing and analysis

The quantity and quality of RNA samples from sorted cells were analyzed using the bioanalyzer form Agilent in combination with the Agilent RNA 6000 Nano. RNA samples with RNA Integrity Number (RIN) > 7 were further used for library preparation using the 3’ mRNA-Seq Library Prep Kit Protocol for Ion Torrent (QuantSeq-LEXOGEN^™^) according to manufacturer’s instructions. The quantity and quality of libraries were assessed using the DNA High Sensitivity Kit in the bioanalyzer, according to the manufacturer’s instructions (Agilent). Libraries were subsequently pooled and templated using the Ion PI IC200 Chef Kit (ThermoFisher Scientific) on an Ion Proton Chef Instrument or Ion One Touch System. Sequencing was performed using the Ion PI^TM^ Sequencing 200 V3 Kit and Ion Proton PI™ V2 chips (ThermoFisher Scientific) on an Ion Proton™ System, according to the manufacturer’s instructions. The RNA-Seq FASTQ files were mapped using TopHat2 [43], with default settings and using additional transcript annotation data for the mm10 genome from Illumina iGenomes (https://support.illumina.com/sequencing/sequencing_software/igenome.html).

According to the Ion Proton manufacturers recommendation, the reads which remained unmapped were submitted to a second round of mapping using Bowtie2 [44] against the mm10 genome with the very-sensitive switch turned on and merged with the initial mappings. Through metaseqr R package [45], GenomicRanges and DESeq were employed in order to summarize bam files of the previous step to read counts table and to perform differential expression analysis (after removing genes that had zero counts over all the RNA-Seq samples).

Downstream bioinformatics analysis and visualization tasks were performed using InteractiveVenn for Venn diagrams (www.interactivenn.net) [46] and the Functional Annotation tool from DAVID for Gene Ontologies (<u>david.ncifcrf.gov</u>) [47, 48]. Volcano plots and heatmaps were generated in R using an in-house developed script utilizing the packages ggplot2, gplots and pheatmap (https://cran.r-project.org/web/packages/pheatmap/index.html) [49–51]. RNA-seq datasets have been deposited in NCBI’s Gene Expression Omnibus [52] and are accessible through the GEO Series accession number GSE117308.

### Comparison with single cell datasets

The positive cluster marker genes of the healthy and DSS treated mouse, from the public dataset GSE114374, were used as gene signatures for each cell type identified in the single cell analysis. For each gene of a signature z-scores of log2 normalized expression values were calculated for the bulk RNA-seq samples GC(GC1, GC2, GC3), TC(TC1, TC2, TC3), UC(UC1, UC2, UC3) and GDS(GDS1, GDS2, GDS3), TDS(TDS1, TDS2, TDS3), UDS(UDS1, UDS2, UDS3). In the boxplots of Figure 1G and 5D the mean z-score of GC1, GC2, GC3 and GDS1, GDS2, GDS3 are displayed respectively.

### Statistical analysis

Data are presented as mean ± SD. Statistical significance was calculated by Student’s t-test or one-way ANOVA for multiple comparisons. The D’Agostino Pearson test was used to test if the dataset followed a normal distribution. Welch’s correction was used for samples that showed unequal variance. P-values ≤ 0.05 were considered significant. Data were analysed using the GraphPad Prism 8 software.

## Acknowledgements

We would like to thank Michalis Meletiou for mouse genotyping. We would also like to thank the Genomics Facility of BSRC “Alexander Fleming”, and specifically Vaggelis Harokopos for performing all RNA sequencing experiments and Martin Reczko and Panagiotis Moulos for initial bioinformatics analyses. We acknowledge support of this work by the InfrafrontierGR Infrastructure, co-funded by Greece and the European Union (European Regional Development Fund), under NSRF 2014-2020, MIS 5002135), which provided mouse hosting and phenotyping facilities, including, histopathology, flow cytometry, and advanced microscopy facilities. This work was supported by a grant from the Stavros Niarchos Foundation to the BSRC “Alexander Fleming” as part of the Foundation’s initiative to support the Greek research center ecosystem, a grant from the Hellenic Foundation for Research & Innovation (H.F.R.I.) to MES (Grant No 1687), the FP7 Advanced ERC grant MCs-inTEST (Grant Agreement No 340217) to GK, a grant from the European Crohn’s and Colitis Organisation to VK and a grant from the Fondation Sante to VK.

